# Evaluation of the nano Liquid Chromatography coupled with Zeno TOF 7600 Mass Spectrometer for Cell Type-Resolved Spatial Proteomics

**DOI:** 10.1101/2025.06.23.660827

**Authors:** Keren Zhang, Xi Wang, Wan He, Jiani Wang, An He, Binbin Zheng, Yangqiu Li, Ruilian Xu, Weina Gao, Henry Lam, Ruijun Tian

## Abstract

In this study, we systematically evaluated the nanoLC-Zeno TOF platform using data-independent acquisition (DIA) and identified 3,142 to 7,059 protein groups from 1 to 200 ng of standard K562 digest. Additionally, a targeted acquisition strategy detected 184 peptides with enhanced signal using Zeno activation. To achieve spatial cell-type resolved proteomics, we developed a workflow integrating image-guided laser capture microdissection for analysis of gastric cancer tissue. A 30-minute nanoLC gradient facilitated the quantification of 2,514 proteins from 0.2 mm² regions representing cancer cells, cancer-associated fibroblasts (CAFs), and immune cells, as defined by multiplex immunohistochemistry. This spatial proteomics approach revealed cellular heterogeneity within the tumor microenvironment of a clinical gastric cancer sample, and targeted validation identified differential expression of 12 cancer-associated proteins comprising 103 peptides between cancer cell-enriched and CAFs-enriched regions. This workflow offers a practical strategy for spatial visual proteomic analysis in clinical tissue samples.

## Introduction

Spatial proteomics has received much attention in recent years (1,2) due to its ability to reveal proteome heterogeneity in tissue samples (3), particularly in complex tumor microenvironments where cell type-specific protein signatures are essential for understanding disease mechanisms (4). Additionally, liquid chromatography coupled with mass spectrometry (LC-MS/MS) have enabled several spatial cell type-specific analyses, such as Deep Visual Proteomics (5) and Spatial and Cell-type Proteomics (SCPro) workflow (6) by integrating with multiplex immunohistochemistry (mIHC) and laser capture microdissection (LCM). These approaches predominantly rely on nano-flow liquid chromatography (nanoLC) with advanced instrumentation design to achieve sensitive proteomic detection from limited tissue samples.

The Zeno TOF 7600 mass spectrometer incorporates a Zeno trap mechanism designed to improve ion transmission efficiency, achieving nearly 100% ion utilization through Zeno trap activation, which substantially enhances MS/MS sensitivity (7). Studies using SWATH data-independent acquisition (DIA) have demonstrated substantial gains in proteome coverage and sensitivity with this platform. For instance, analytical-flow LC coupled with Zeno SWATH has enabled biomarker discovery and pathway analysis in human and cynomolgus monkey tissues (8), Similarly, micro-flow LC setups have achieved identification of over 5,000 protein groups from as little as 62.5 ng of K562 digest (9). Gu et al. demonstrated that combining narrow isolation windows with Zeno SWATH acquisition on the Zeno TOF 7600 significantly improves proteomic identification depth and quantification precision from complex biological samples compared to conventional SWATH approaches (10). While these investigations have demonstrated the potential of the Zeno TOF 7600 platform using analytical- and micro-flow LC, they have not evaluated its performance coupled with nanoLC, which remains the predominant approach for high-sensitive proteomic analysis. Given that Zeno SWATH DIA has already been shown to provide deep proteome coverage, the combination of nanoLC with the Zeno TOF 7600 is expected to deliver optimal performance for spatial proteomic analyses with the possibility that has yet to be systematically explored.

For applications in tissue samples, DIA-MS on Zeno TOF 7600 system was applied to a mouse model of acute kidney injury (11). By generating a comprehensive kidney sample specific deep spectral library with 120-minute gradient separations, researchers achieved identification of 3,945 protein groups with more than half of the quantified proteins exhibiting significant alterations. More recently, Humphres et al. (12) suggest that the performance gap between nanoflow- and microflow-LC has become minimal for many proteomic applications. They optimized a workflow with a 5-minute trap, 30-minute LC gradient, 50 variable SWATH windows, and a 10 ms MS2 accumulation time. This allowed sample loads as low as 400 ng, outperformed conventional 90-minute gradients on the TripleTOF 6600, and enabled identification of ∼2,000 protein groups from eight rat organs. In their subsequent work (13), analysis of matched formalin-fixed paraffin-embedded (FFPE) and fresh-frozen (FF) rat tissues from four organs demonstrated that, using the same workflow with 200 ng peptide input, testis samples yielded the highest protein groups identification number of 2,403 in FFPE and 3,214 in FF.

In summary, we aimed to systematically evaluate the Zeno TOF 7600 coupled with a nanoLC platform for both profiling and targeted proteomics, thereby providing a comprehensive assessment of its performance. By further integrating with our previously reported SCPro workflow (6) for spatial visual proteomic analysis, we established a cell-type resolved spatial proteomics approach optimized for the Zeno TOF 7600 platform and applied to clinical gastric cancer tissue samples. Through these studies, we seek to demonstrate the broader applicability of this platform in spatial visual proteomics and to facilitate its adoption in diverse biological and clinical contexts.

## Experimental Procedures

### Standard Samples and Tissue Samples

For calibration and quality control purposes, we utilized standard peptide mixtures including MS Synthetic Peptide Calibration Kit PepCalMix (SCIEX, P/N 5045759) and iRT (Biognosys, Ki-3002-1). Custom synthetic peptides were acquired from GeneScript for targeted analyses. Method evaluation was conducted using the SWATH® Acquisition Performance Kit containing K562 cell digest (SCIEX, P/N 5045757). Additionally, laboratory-cultured samples including HeLa cells, Yeast, and E. coli were prepared to assess versatility across diverse biological matrices. Protein extraction and enzymatic digestion of these cultured samples were performed following the Filter-Aided Sample Preparation (FASP) protocol (14). All processed samples were subsequently subjected to LC-MS/MS analysis to comprehensively evaluate the performance characteristics of our experimental workflows.

### Spatial Sample Preparation and Multiplex Immunohistochemistry (mIHC) Staining

Formalin-fixed, paraffin-embedded (FFPE) gastric cancer tissue sections with a thickness of 4 μm were used in this study. The sections were mounted on membrane slides and subjected to 4-color mIHC using tyramide signal amplification (TSA) technology. Staining was performed using an automated staining machine (Dartmon, AS330-PLUS), which completed four rounds of staining to label primary antibody PanCK (CST#4545, 1:1000), αSMA (Proteintech#14395-1-AP, 1:2000), and CD45 (CST#70257, 1:500). Antigen retrieval was conducted using heat-induced epitope retrieval (HIER) in a pH 6.0 buffer at 100°C for 20 minutes in each round. Primary antibodies were incubated for 60 minutes, followed by secondary antibody incubation for 30–45 minutes, and fluorescent dye staining was performed for 10 minutes in specific channels (520 nm for PanCK, 570 nm for αSMA, and 650 nm for CD45). Finally, DAPI staining was applied to all sections for nuclear visualization by incubating the slides in the dark at room temperature for 10 minutes, followed by mounting. The automated staining process ensured consistent and reproducible labeling across all samples.

### Imaging and Analysis

Tissue imaging was performed using the TissueFAXS imaging platform equipped with a 40X objective, achieving a resolution of 0.16 μm/pixel. Image analysis was conducted using TissueGnostics StrataQuest v7.1 software. A random forest algorithm was used to train the classifier for accurate identification of target features. Cancer cells, CAFs, immune cells were analyzed by applying specific gating thresholds to exclude false-positive cells and excessively small cells. To preserve intact cell morphology for subsequent laser capture microdissection (LCM), the mask was expanded by 15 pixels (0.16 μm/pixel) during the analysis process. VistaNavi software (BayOmics, version 1.0.2) was used to process scanned images, mask images, and XML files containing positional information (typically represented as rectangular frame paths from the LCM system). The software generated executable cutting-path XML files with adjusted scale and orientation. LCM was performed on a CellCut system (Molecular Machines & Industries, MMI) with a 20X objective. The laser parameters were set as follows: velocity: 40 μm/sec, focus: 164 μm, power: 30%. Each dissection was repeated once.

### Sample preparation

All LCM samples were processed using the SISPROT PAC kit (BayOmics#J18-01-000041) according to the manufacturer’s protocol. The resulting peptides were lyophilized and reconstituted in 0.1% formic acid water prior to LC-MS/MS analysis.

### LC-MS/MS

Nano-flow liquid chromatography (LC) analyses were performed using a M-Class UPLC system (Waters) operated in direct-inject mode. The mobile phases consisted of (A) 0.1% (v/v) formic acid in 98:2 (v/v) water/acetonitrile and (B) 0.1% (v/v) formic acid in 98:2 (v/v) acetonitrile/water. Chromatographic separations were conducted on a home-made column (75 μm I.D. × 15 cm, 1.9 μm C18) at a flow rate of 300 nL/min. Gradient elution protocols were applied with a mobile phase B range of 3–35%. For K562 dilutions, samples at concentrations of 200 ng/μL and lower were analyzed using gradient durations of 10, 20, 30, 45, or 60 minutes, with a 1 μL injection volume. ZT18 mixtures were analyzed under identical chromatographic conditions, employing a fixed 30-minute gradient (3–35% B) and a 1 μL injection volume. Gastric cancer tissue samples were processed similarly but with a 3 μL injection volume and a 30-minute gradient (3–35% B). These conditions ensured consistent chromatographic resolution across all sample types. Data-independent acquisition (DIA) experiments were conducted using a Zeno TOF 7600+ hybrid quadrupole time-of-flight mass spectrometer (SCIEX) equipped with a horizontal nanoflow electrospray ionization (NSI) source Turbo V and OptiFlow interface. Standardized ion source parameters were applied across all experiments, including an ion spray voltage of 3800 V, curtain gas at 35 psi, nebulizing gas (Gas 1) at 18 psi, and an interface heater temperature of 175°C. Three Zeno SWATH DIA acquisition methods were employed, each utilizing a Q1 mass range of 400–900 Da with differential number of variable-width windows and MS/MS accumulation times:

(1) 38 windows with a 32 ms accumulation time, (2) 65 windows with a 20 ms accumulation time, and (3) 85 windows with an 18 ms accumulation time. All methods incorporated a TOF-MS precursor survey scan (400–1500 Da) with a 200 ms accumulation time, collision-induced dissociation (CID) with dynamic collision energy ramping, and Zeno trap pulsing. The MS/MS mass range spanned 140–1750 Da. Triplicate acquisitions were performed for each experimental condition to ensure reproducibility. Additionally, ZT Scan DIA experiments were optimized for 30-minute chromatographic gradients, using a peak width at half-height (PWHH) setting of >4.5 seconds. Data was acquired in triplicate for all samples under these conditions. The targeted mode was configured in MRM^HR^, using a 250 ms full-scan duration with the same scan range and source parameters as those set in the SWATH mode. Each MS2 scan was executed with an acquisition time of 15 ms, employing an optimized collision energy (CE) for enhanced fragmentation efficiency.

## Data Processing

For method evaluation by cell digest sample, MS data was processed using DIA-NN (15) software (version 1.8.16, https://github.com/vdemichev/DiaNN) against a pre-constructed spectral library comprising K562 and HeLa cell line proteomes (SCIEX). Key search parameters included a precursor m/z range of 400–900, a fragment m/z range of 140–1750, and a mass accuracy tolerance of 20 ppm for precursors and 0 ppm for MS1. For ZT Scan DIA datasets, the --scanning-swath command was applied to optimize processing. Triplicate files from identical loading and gradient conditions were processed independently to ensure statistical robustness and minimize variability. For spatial proteomic data analysis, raw mass spectrometry data were processed using Spectronaut 18.2.2 software, directDIA mode with default setting under database UP000005640_9606_HUMAN (20406 entries). For targeted proteomic data, processing was performed using Skyline 23.1.0 with default peptide settings. The ion match tolerance was set to 0.2 m/z, allowing only precursor ions with charge states of +2 and +3, and fragment ions (b and y types) with a charge state of +1. Monoisotopic precursor and product ion masses were utilized, with collision energy parameters optimized for SCIEX instrumentation. MS1 filtering was set to "None" and the MS/MS acquisition method was configured to PRM, employing a resolving power of 30,000 through the TOF analyzer. All matching scans were included.

### Statistics Analysis

Proteins were filtered to retain only those with at least three valid values in at least one experimental group. The raw intensity values were log2-transformed, and any NA values were replaced with zeros. Differentially expressed proteins between two groups were presented by scatter plot analysis, with a fold-change threshold of 4 and label protein name with Log2 ratio >= cutoff and p adjust value <= 0.05. Differential expression analysis among three groups was performed using One-way ANOVA with Benjamini-Hochberg correction for multiple testing was performed on the filtered dataset, and proteins with adjusted p-values ≤ 0.01 were subjected to hierarchical clustering using Pearson correlation distance and average linkage using a hierarchical clustering heatmap. Principal component analysis (PCA) was applied to assess variance and clustering among the studied groups. Gene Ontology (GO) enrichment analysis was performed using Metascape (16).

### Ethics Approval and Patient Consent Statement

Gastric cancer tissue sections were collected following the ethical approval (SYL-202124-02). All experiments were performed in accordance with the Declaration of Helsinki.

## Result

### Performance of Data Independent Acquisition with Zeno SWATH and ZT Scan

Prior studies have shown that the implementation of SWATH DIA with the Zeno On mode demonstrated significant improvements in data acquisition performance compared to Zeno Off (1–2). We first evaluate the fragment ions using an established quality assessment method (6), SWATH analysis of a 200 ng standard K562 digest sample showed that Zeno On mode increased both fragment ion acquisition per cycle and MS2 peak intensities compared to Zeno Off (Fig. 1A-B). Furthermore, despite acquiring a lower number of MS2 scans, Zeno SWATH demonstrated a higher spectral matching rate compared to conventional SWATH acquisition (Fig. 1C). This indicates that the Zeno SWATH approach more effectively utilizes the acquired spectra to sample a broader diversity of precursor ions, thereby enhancing the overall peptide and protein identification efficiency. Consequently, Zeno SWATH identified >30% more protein groups and >50% more peptides than standard SWATH across 20-200 ng K562 digest samples (Fig. 1D).

**FIG. 1.**
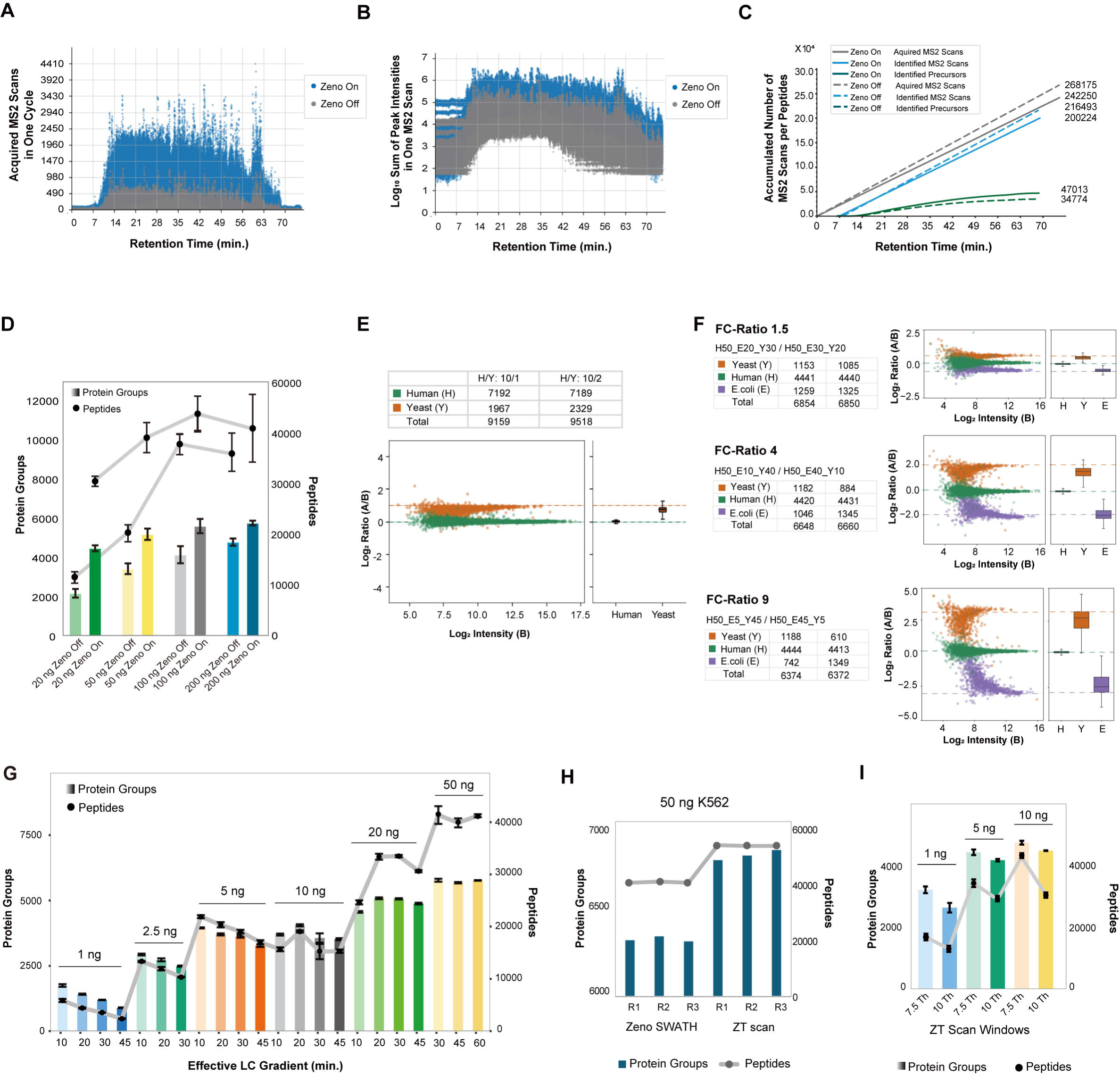
Performance SWATH with Zeno Off and Zeno On Mode. **(A)** Number of MS2 scans over LC gradient in SWATH with Zeno Off (Grey) and Zeno On (Blue). **(B)** MS2 intensities for each scan over LC gradient in SWATH with Zeno Off (Grey) and Zeno On (Blue). **(C)** Accumulation number of acquired and identified MS2 scan with identified Precursors. **(C)** Number of identifications of 20 ng, 50 ng, 100 ng and 200 ng K562 sample injections with 3 species using SWATH with Zeno Off and Zeno On mode. **(E)** Protein identification and quantification in a human-yeast mixture at 1/10 and 2/10 ratios, with a total of 200 ng per sample and three replicates. Each point represents an individual protein, and the boxplot shows the distribution of logD ratio values for human (green) and yeast (orange). **(F)** Protein identification and quantification in a three-species mixture (human, yeast, and E. coli) at six different ratios, with a total of 100 ng per sample and three replicates. Ratios of 1.5, 4, and 9 are shown for each species. Each point represents an individual protein, and the boxplot shows the distribution of logD ratio values for human (green), yeast (orange), and *E. coli* (purple). **(G)** Number of identifications from 1 ng to 50 ng K562 samples with three replicates, using Zeno SWATH under different effective LC gradients. **(H)** Number of identifications for 50 ng K562 samples using Zeno SWATH and ZT Scan. **(I)** Number of identifications from 1 ng to 10 ng K562 samples using ZT Scan with 7.5 Th and 10 Th scanning modes.

To assess Zeno SWATH quantification accuracy, we analyzed 200 ng samples of HeLa and yeast protein digests mixed at 1:10 and 1:5 ratios. Zeno SWATH accurately quantified 1,967 and 2,329 yeast protein groups respectively in these mixtures where yeast represented only a minor fraction of human proteins (Fig. 1E). This precision was achieved while maintaining extensive proteome coverage, with 9,518 total protein groups identified. To validate quantitative performance with more complex samples, we analyzed six multi-species samples containing defined proportions of digested HeLa, yeast, and *E. coli* (17). Each sample contained a total protein amount of 100 ng and was acquired using Zeno SWATH for relative quantification across species. The distribution of protein quantification ratios (Fig. 1F) closely mirrored with the expected mixing proportions across all samples. To assess Zeno SWATH performance with limited sample input (18), we evaluated the effect of shortened chromatographic gradients on protein and peptide identifications (Fig. 1G). we evaluated shortened chromatographic gradients on identification rates (Fig. 1G). For samples below 5 ng, shortened gradients significantly improved protein and peptide identifications. Using a 10-minute effective gradient, we identified 1,820 protein groups and 6,146 peptides from 1 ng K562 digest and 3,989 protein groups and 22,342 peptides from 5 ng input. Notably, our 2.5 ng results yielding 2,999 protein groups and 13,596 peptides surpass leading microflow-LC reports(9), likely due to the sensitivity of nano-LC at low sample inputs.

Additionally, we compared ZT Scan, also known as scanning SWATH (19) , with Zeno SWATH using replicate analyses of 50 ng K562 digest (Fig. 1H). ZT Scan showed improved reproducibility with more proteins exhibiting low variability across replicates. Identified protein groups increased from 6,491 to 7,141 and peptide identifications from 44,290 to 61,026. The percentage of proteins with CV values below 20% and 10% improved by 24.2% and 45.7% respectively, while peptide improvements were 55.8% and 86.3% (Supplemental Fig. 1). When comparing 7.5 Th and 10 Th scanning windows under a 30-minute LC gradient (Fig. 1I), the 7.5 Th window consistently yielded more identifications across all sample inputs. With 1 ng of K562 digest, the 7.5 Th mode identified 3,142 protein groups and 17,367 peptides, comparable to Zeno SWATH results with 5 ng input under identical conditions. The improvement became less pronounced at higher sample inputs (5-50 ng), suggesting effectiveness under low-input conditions.

### Targeted Proteomics Evaluation using Zeno MRM^HR^

We evaluated the performance for targeted proteomics firstly using iRT peptide mixture (11 peptides) and PepCalMix (20 heavy isotope-labeled peptides). In our MRM^HR^ method, we allocated 250 ms for MS1 full scan and 10 ms per peptide for MS2 scan. This 10 ms accumulation time produced well-defined peaks, with minimal signal improvement when extended to 100 ms (Supplemental Fig. S2A). The Zeno On mode significantly enhanced peptide signal intensity. All 11 iRT peptides across various retention times and concentration ranges showed >5-fold signal enhancement under Zeno On compared to Zeno Off conditions (Fig. 2A-B). This improvement results from Zenotrap efficient ion accumulation and release system, which achieves near-complete ion detection versus approximately 30% efficiency in Zeno Off mode. Individual fragment ions in Zeno Off mode displayed signal intensities below 1,000, potentially compromising detection in complex samples due to background interference. Conversely, Zeno On mode elevated these signals to over 7,000, substantially improving fragment ion matching and enabling more sensitive quantification potential (Fig. 2C).

**FIG. 2.**
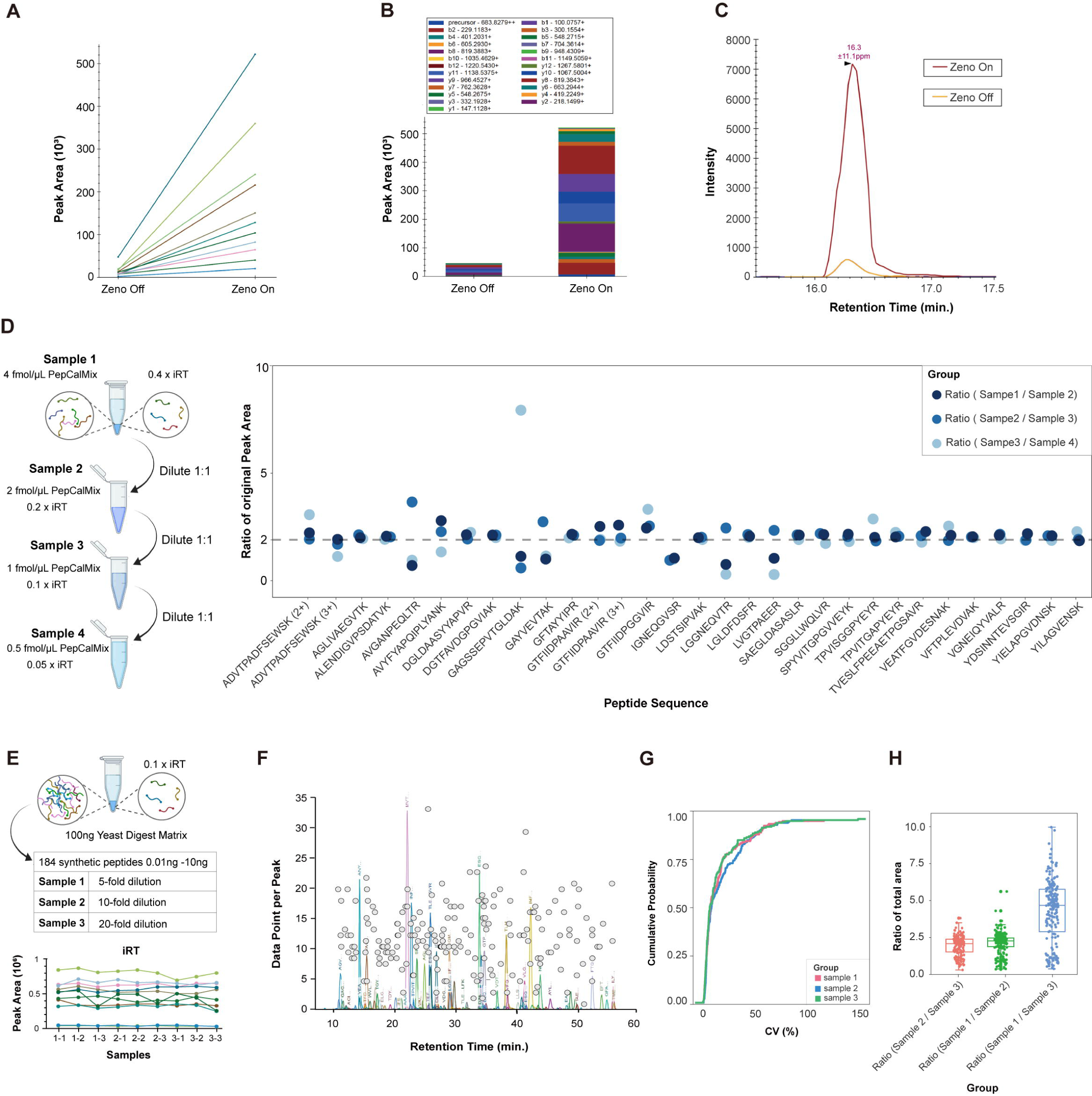
Performance of MRM^HR^ with Zeno Off and Zeno On Mode. **(A)** Peak area of 11 peptides from iRT detected on MRM^HR^. Each line demonstrates increases in each peak area from Zeno Off to Zeno On. **(B)** Peak area of a peptide accumulated with all fragment ions under Zeno Off and Zeno On. **(C)** Intensity comparison of a fragment ion detected on MRM^HR^ with Zeno Off to Zeno On. **(D)** Sample setting by gradient dilution of iRT and PepCalMix standard peptide mixtures; Quantitative ratios at three concentration levels. **(E)** Sample setting of 184 synthetic peptides in yeast digest matrix with equal amount of 0.1 x iRT for each diluted sample; iRT peptide peak areas across three groups with three replicates shown below. **(F)** The number of data point per peak of each precursor over LC gradient. **(G)** CV (%) of total detected peptides in three different concentration groups. **(H)** The distribution of total area ratios among 3 groups in different concentrations.

Next, to assess quantitative accuracy, a mixture of iRT and PepCalMix peptides was prepared starting at 0.4 x and 4 fmol/μL respectively and subjected to 1:1 serial dilution to generate four concentration points (Fig. 2D). Pairwise comparisons across 31 peptides revealed that most peptides closely aligned with the expected 2-fold variation between sample groups. Peptide ADVTPADFSEWSK exhibited deviation from the expected ratio in low-concentration samples, with divergent behavior between its 2+ and 3+ charge states, while maintaining expected ratios in medium-high concentration comparisons. Peptide DGTFAVDGPGVIAK demonstrated consistently accurate abundance ratios across all groups with high-quality peak extraction (Supplemental Fig. S2D). Conversely, GAGSSEPVTGLDAK showed a significant outlier in the low-concentration group due to sample matrix interference, and ratio compression in higher concentration groups attributed to peak missing in one replicate. Hydrophilic peptides eluting early in the gradient showed increased susceptibility to loss, potentially due to adsorption effects amplified by the three-day interval between replicate acquisitions.

One of the primary challenges in implementing routine targeted proteomics lies in the complexity of the sample matrix, limitations in dynamic range, and the number of target peptides, which complicates the development of optimized methods for quantifying numerous peptides simultaneously. To address this, we utilized 184 synthetic peptides ranging from 0.01 to 10 ng prepared in three dilutions of 5-fold, 10-fold, and 20-fold spiked into 100 ng Yeast digest with 0.1× iRT peptides as equal-amount quality control. Results showed consistent detection of iRT peptides across all samples and replicates as illustrated in Figure 2E. Using a 60-minute nano LC gradient with unscheduled target MS, we detected 171 peptides achieving a 93% detection rate, with sufficient data points per peak average over 10 (Fig. 2F). Over 70% of the detected 171 peptides in three replicates in each sample achieved quantification coefficient of variation (CV) below 20% (Fig. 2G). Relative quantification between sample groups revealed median values aligned with expected ratios as demonstrated in Figure 2H, indicating accurate peptide quantification despite complex matrix background. These results validate our targeted proteomics workflow for reliable quantification of multiplex targets with wide dynamic range.

### Implementing Spatial Proteomics using Zeno TOF 7600

Building on the demonstrated capabilities of the nano-LC coupled Zeno TOF 7600 system for both deep proteomic profiling and targeted multiplexed analysis, we extended its application to a spatial proteomics workflows using FFPE clinical gastric cancer samples. Tissue sections were processed using LCM to isolate specific regions of interest followed by SISPROT sample preparation (Fig. 3A). Histological and molecular characterization was performed using hematoxylin and eosin (H&E) staining and mIHC. LCM was employed to isolate specific tissue regions of interest, followed by sample preparation using the SISPROT to generate peptides.

**FIG. 3.**
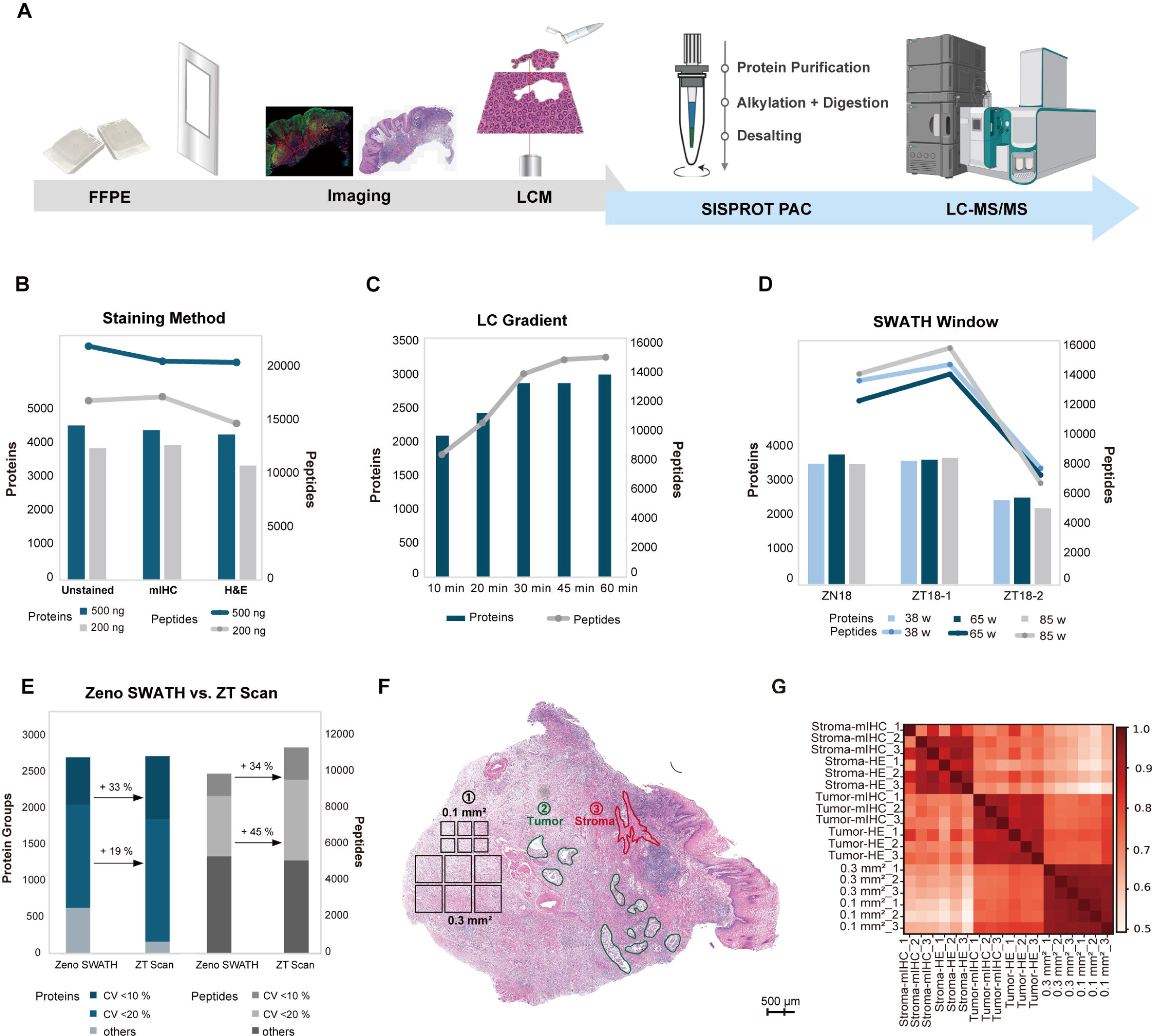
Improvement of spatial proteomics workflow for FFPE tissue samples. **(A)** Workflow of spatial proteomics analysis for FFPE tissue samples. Tissue sections were imaged for histological or immunophenotypic features, followed by laser capture microdissection (LCM) to isolate specific areas of interest, followed by SISPROT sample preparation and analyzed using LC-MS/MS. **(B)** Comparison of protein and peptide identification numbers at 200 ng and 500 ng from LCM tissue samples processed under three conditions: unstained, mIHC-stained, and H&E-stained. **(C)** LC gradient optimization using 0.2 mm² ZT18-2 LCM samples. Each data corresponds to a 0.2 mm² sample with mixed samples derived by aggregating multiple areas for analysis. **(D)** Evaluation of protein and peptide identification numbers under different SWATH window settings (38, 65, and 85 windows) on 0.2 mm² LCM samples with 3 sample types. **(E)** Comparison of peptide identification numbers of ZT18-2 acquired by Zeno SWATH and ZT Scan under CV < 20% (Darker blue) and CV < 10% (Grey). **(F)** Overview of H&E-stained ZT18-2 and ROI (Region of Interest) of each cell type in black lines. 12 square area: 0.1 mm² and 0.3 mm², 12 Tumor-enriched region, 12 Stroma-enriched region, with collection area of 0.1 mm² for each sample. **(G)** Correlations among 3 types of samples stained H&E and mIHC, with 3 replicates of each stained sample. The color gradient (light to dark red) represents correlation coefficients from 0.5 to 1.0.

Next, we went on to investigate different parameters that affect the sensitivity of spatial proteomic analysis. Comparative analysis for staining method revealed that both H&E and mIHC staining methods yielded comparable protein and peptide identification numbers to unstained sections, which identified 3,705 and 4,331 protein groups and 17,567 and 22,691 peptides at 200 ng and 500 ng injection levels, respectively (Fig. 3B). This demonstrates that staining does not significantly compromise protein identification. We then optimized the workflow using a tissue area of 0.2 mm² to reflect realistic sample limitations in clinical settings. Our gradient optimization showed that a 30-minute effective gradient produced 2,859 protein groups and 13,688 peptides, comparable to the 60-minute gradient with only a 10% reduction in peptide yield while improving time efficiency (Fig. 3C). Evaluation of SWATH acquisition window settings across three distinct tissue regions demonstrated minimal differences in protein identification numbers between 38, 65, and 85 variable windows (20,21) (Fig. 3D), indicating marginal effect of window configuration on identification performance. Furthermore, ZT Scan for a 0.2 mm^2^ sample revealed no significant difference in protein identification numbers of 2722 compared to Zeno SWATH of 2707, but showed more stable CVs, especially in peptides identification (Fig. 3E), making it the preferred mode for consistent and reliable spatial proteomics analysis.

Finally, to evaluate spatial proteomic differences, tumor-rich and stroma-rich regions were manually identified using H&E staining (Fig. 3F) and mIHC (Supplemental Fig. S3B) and compared 0.1 mm² and 0.3 mm² sampling areas. Analysis revealed significant correlation differences between tumor-rich and stroma-rich regions, confirming their distinct proteomic profiles. Conversely, the 0.1 mm² and 0.3 mm² areas showed high correlation, reflecting their relative homogeneity (Fig. 3G). Strong correlations were observed between mIHC fluorescence and H&E staining data from identical sample types (Supplemental Fig. S3C). These correlation patterns highlight the necessity of cell type-based spatial proteomics analysis, as they demonstrate that proteomic composition is primarily determined by cell type rather than sampling area size. The distinct proteomic signatures between tumor and stromal regions, coupled with the consistency within each cell type, underscore the importance of cell type-specific analysis for accurate characterization of tumor microenvironments.

### Image-Guide Cell Type-Resolved Segmentation and Dissertation

For cell-type-resolved spatial proteomics sample collection, we performed four-color mIHC staining using DAPI, pan-CK, α-SMA and CD45 to label the cell nuclei, cancer cells, cancer-associated fibroblasts (CAFs) and immune cells, respectively. Panoramic screening of whole-slide images identified a tissue ROI of 8.43 mm² containing all cancer cells (Fig. 4A). Using the Classifier module, we labeled representative cancer cells as positive examples and other regions as negative examples to train a random forest model that generated cancer cells shape masks (Fig. 4B). We refined the segmentation by excluding DAPI-negative cells, areas smaller than 1500 μm², and cells with marker signal intensities below 100 arbitrary units. Histogram-based gating selected tumor regions within specific size ranges, yielding 45 tumor-positive shapes (Fig. 4C). The finalized masks were processed with VistaNavi software to generate cutting-path XML files for the LCM system, with paths manually adjusted to ensure precise alignment with tissue morphology (Fig. 4D). Post-dissection fluorescence microscopy confirmed the integrity and specificity of collected samples on a sticky cap (Fig. 4E). In total, three replicate tissue slides, each containing three distinct cell types with a collection area of 0.2 mm² per type, were successfully obtained for subsequent proteomic analysis.

**FIG. 4.**
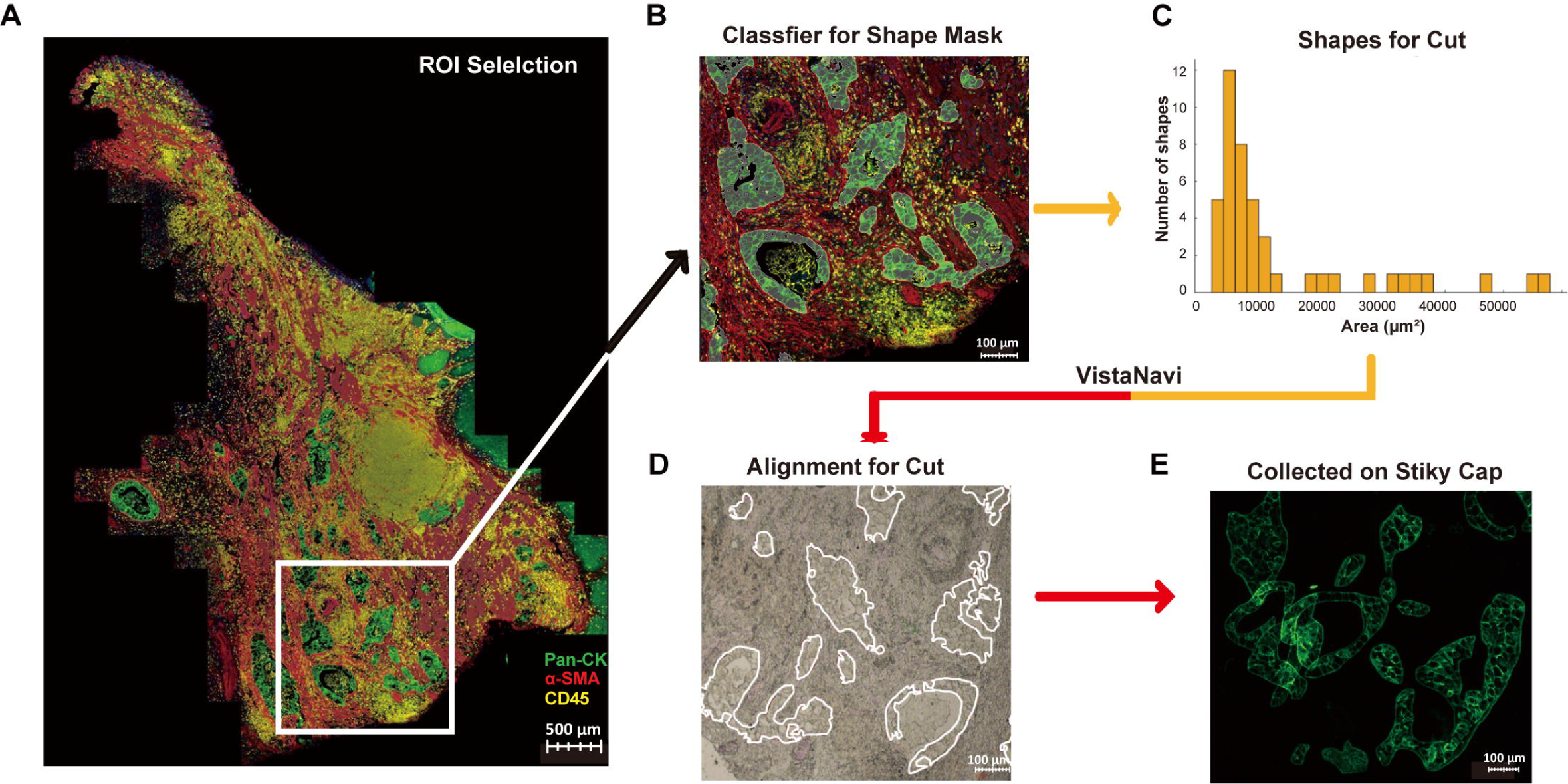
Cell type-resolved spatial proteomics sample collection based on VistaNavi. (A) ROI selection for the ZT18-2 sample, encompassing all tumor regions based on mIHC staining, cancer cell marker Pan-CK (green), α-SMA for CAFs (red), CD45 for immune cells (yellow) and DAPI (blue) for nucleus. (B) Cancer cell mask (white outline) generated using a trained random forest classifier and iteratively refined to remove misclassified regions. (C) Histogram-based gating to select cancer cells within biologically relevant size ranges. (D) Alignment of the XML cutting path (white lines) generated by VistaNavi software with real-time tissue morphology on the laser capture microdissection platform. (E) Fluorescence validation of the collected cancer cell shapes on a sticky cap.

### Cell Type-Resolved Spatial Proteomics for a Clinical Gastric Cancer Patient

Following our spatial visual proteomics workflow, we collected 0.2 mm² tissue samples from cancer cells, CAFs, and immune cells. Microdissected samples were prepared using SISPROT PAC, specifically iPAC (6), which effectively cleaned spatial membrane-based LCM tissue samples. We quantified 2,514 protein groups across three cell types, each with three biological replicates (Fig. 5A). Proteomic analysis of cancer samples revealed a dynamic range of protein abundance (Fig. 5B). In which, KRT8 (22), a recognized gastric cancer marker, displayed significantly a high expression level. VIM, known as a surface marker of circulating tumor in gastric cancer (23), was also readily detected. Notably, TGFB1, a protein associated with gastric cancer (24) and chemotherapy response (25) in spatial omics studies, was detected at relatively low abundance. And the highest TGFB1 expression among three cell types was in CAFs samples (Supplemental Fig. 5A), consistent with its primary activity being concentrated in CAF-enriched stromal regions (26). The detection of TGFB1 demonstrates the sensitivity and precision of our technology in capturing heterogeneous expression patterns of key regulatory proteins. Principal component analysis (PCA) showed clear separation among cancer cells, CAFs, and immune cells (Fig. 5C), indicating distinct proteomic profiles for each cell type and confirming the reliability of our spatial segmentation and cell-specific sampling process. A comparative analysis of differentially expressed proteins (DEPs) between cancer cells and CAFs, as well as between cancer cells and immune cells, revealed distinct expression profiles highlighting cancer-specific protein alterations (Fig. 5D). Proteins significantly upregulated in cancer regions compared to both CAFs and immune cells (Both Up, marked in red) included established cancer markers such as KRT8, KRT18, KRT19, S100P, and ANXA4, as well as GOLM1, ACSL6, ASS1, and SLC25A1.

**FIG. 5.**
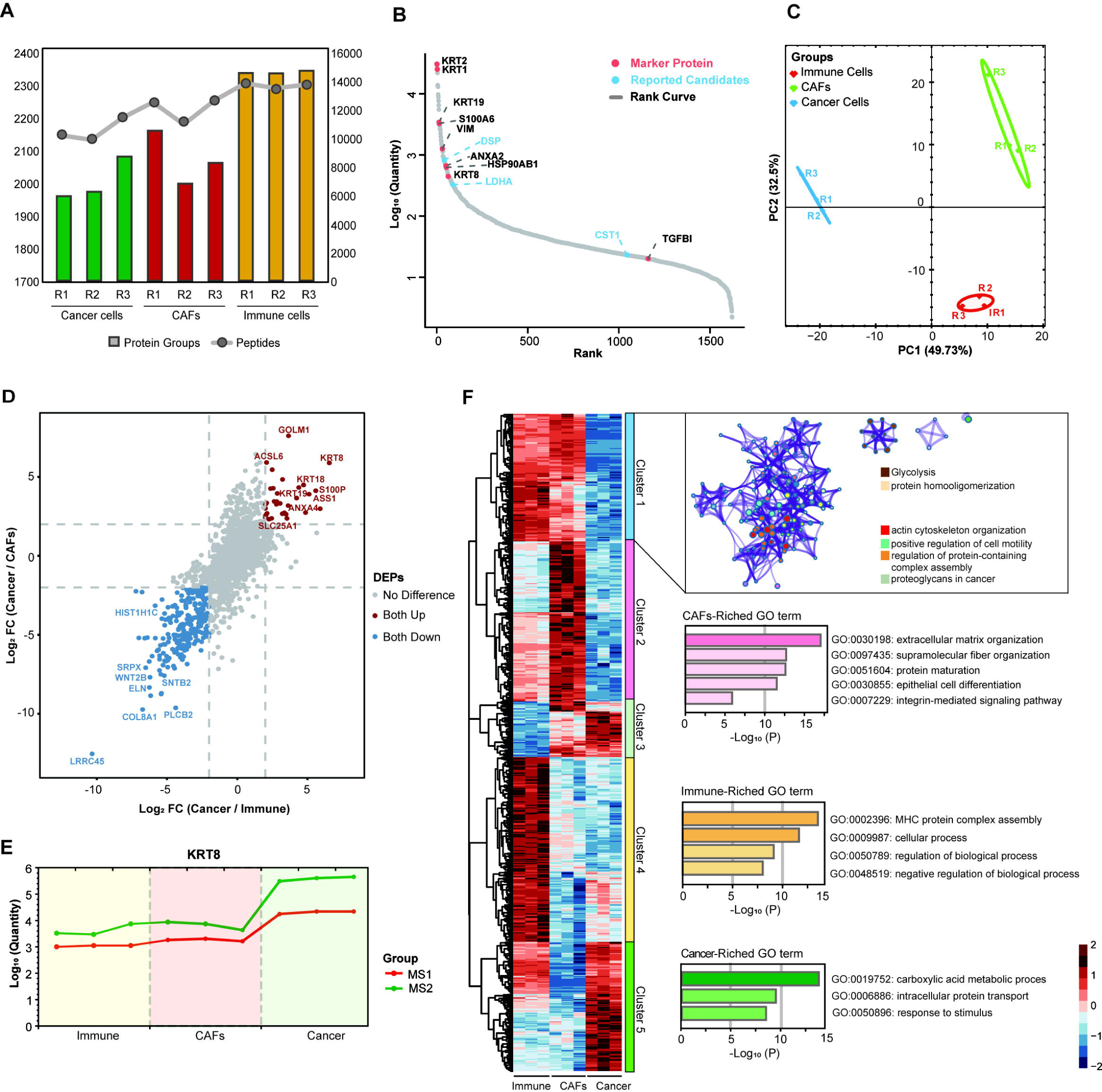
Cell-type spatial proteomics. **(A)** Identification number of three cell-type LCM samples with three replicates. **(B)** Cancer cell protein rank. Marker proteins (red) and proteins reported in the literature to be associated with gastric cancer (blue) are marked. **(C)** PCA plot of three groups: Immune cells (red), CAFs (green), Cancer cells (in blue). PC1 (49.73%) and PC2, (32.5%) define the principal component space. **(D)** Scatter plot for differential protein expression between two comparisons, Cancer vs Immune cell type samples and Cancer vs CAFs with log_2_ (fold change). **(E)** Gastric cancer marker KRT8 Quantity in MS1 and MS2. **(F)** Heatmap generated by one-way ANOVA followed by post-hoc tests. Each cluster was subjected to GO term enrichment analysis. Cluster 1 displays pathways and interaction networks. Clusters 2–5 show pathway enrichment and significance, except Cluster 3, which is not shown.

Conversely, proteins downregulated in cancer regions (Both Down, shown in blue) included LRRC45, COL8A1, PLCB2, ELN, WNT2B, SRPX, HIST1H1C, and SNTB2. These findings identify proteins with cancer-specific expression patterns that potentially define cellular identities within the tumor microenvironment. The gastric cancer marker KRT8 demonstrated consistently high expression in cancer samples using both MS1- and MS2-based quantification methods (Fig. 5E). Between immune cells and CAFs, 513 proteins were differentially expressed with 211 upregulated in immune cells (Supplemental Fig. 5B), including immune-specific proteins such as CD74, HLA-class I and II proteins, and T-complex protein-1. Histone proteins were also significantly upregulated, likely due to the larger nuclei and higher nuclear content of immune cells.

A heatmap of proteomic data showed clear clustering of the three cell types across five distinct clusters (Fig. 5F). Cluster 1 included proteins upregulated in both CAFs and immune cells but downregulated in cancer cells, with GO term enrichment in actin cytoskeleton organization, cell motility, and protein-complex assembly. Cluster 2 showed significant enrichment of upregulated proteins in CAFs, with functional annotations highlighting ECM organization, supramolecular fiber organization, protein maturation, epithelial cell differentiation, and integrin-mediated signaling. Key ECM-related proteins in this cluster included COL4A, COL6A, COL1A families, ITGB1, FBLN3, and LAMA4, supporting the role of CAFs as stromal regulators. EFEMP1, a significant protein in this cluster, has been implicated in gastric cancer progression (27). Cluster 4 contained proteins enriched in immune cells, including LMP2, a subunit of the immunoproteasome (28), with functions related to MHC protein complex assembly and various cellular regulatory processes. Cluster 5 comprised proteins predominantly expressed in cancer cells, involved in carboxylic acid metabolism, intracellular protein transport, and stimulus response. These findings demonstrate the functional heterogeneity within the gastric cancer microenvironment and highlight the distinct molecular signatures of cancer cells, CAFs, and immune cells. The consistent and biologically relevant protein expression patterns observed across cell types confirm the reliability of our developed system for cell type-specific spatial proteomics.

### Region-Specific Targeted Proteomic Assessment

To evaluate targeted proteomics applicability in spatially resolved clinical samples, we selected a cluster of proteins highly correlated with gastric cancer marker KRT8 (Fig. 6A) as targets. As a result, 117 peptides from twelve proteins were generated from the cell-type proteome data without filtering. Two distinct ROIs were set: a cancer-enriched region with a proportion of 38.1%, and another CAFs-enriched region with 1.3% of cancer cells presence of only (Fig. 6B). The targeted approach successfully detected 88% of peptides (103/117), confirming the reliability of our profiling data. All 14 KRT8 peptides detected by ZT Scan were likewise detected by MRM^HR^, with consistently higher expression in the cancer-enriched sample (Fig. 6C), validating our analytical approach. While proteins like KRT18 were quantified using multiple peptides, others such as SLC25A6 relied on single peptide quantification (DFLAGGIAAAISK). The targeted methodology optimized chromatographic peak quality and area, yielding reproducible signals across samples (Fig. 6D-E), though SLC25A6 showed minimal expression differences between regions. Quantification of all 12 proteins revealed significant differential expression patterns (Fig. 6F). Beyond KRT8, several proteins exhibited substantial enrichment in cancer regions compared to CAFs enriched regions: ASS1 (4.3-fold), KRT18 (2.3-fold), TXRD1 (3.0-fold), ANXA4 (2.1-fold), and ALDH2 (2.2-fold). These results demonstrate the efficacy of targeted proteomic approaches for spatial samples and their potential utility for precise quantitative analysis in specialized clinical applications.

**FIG. 6.**
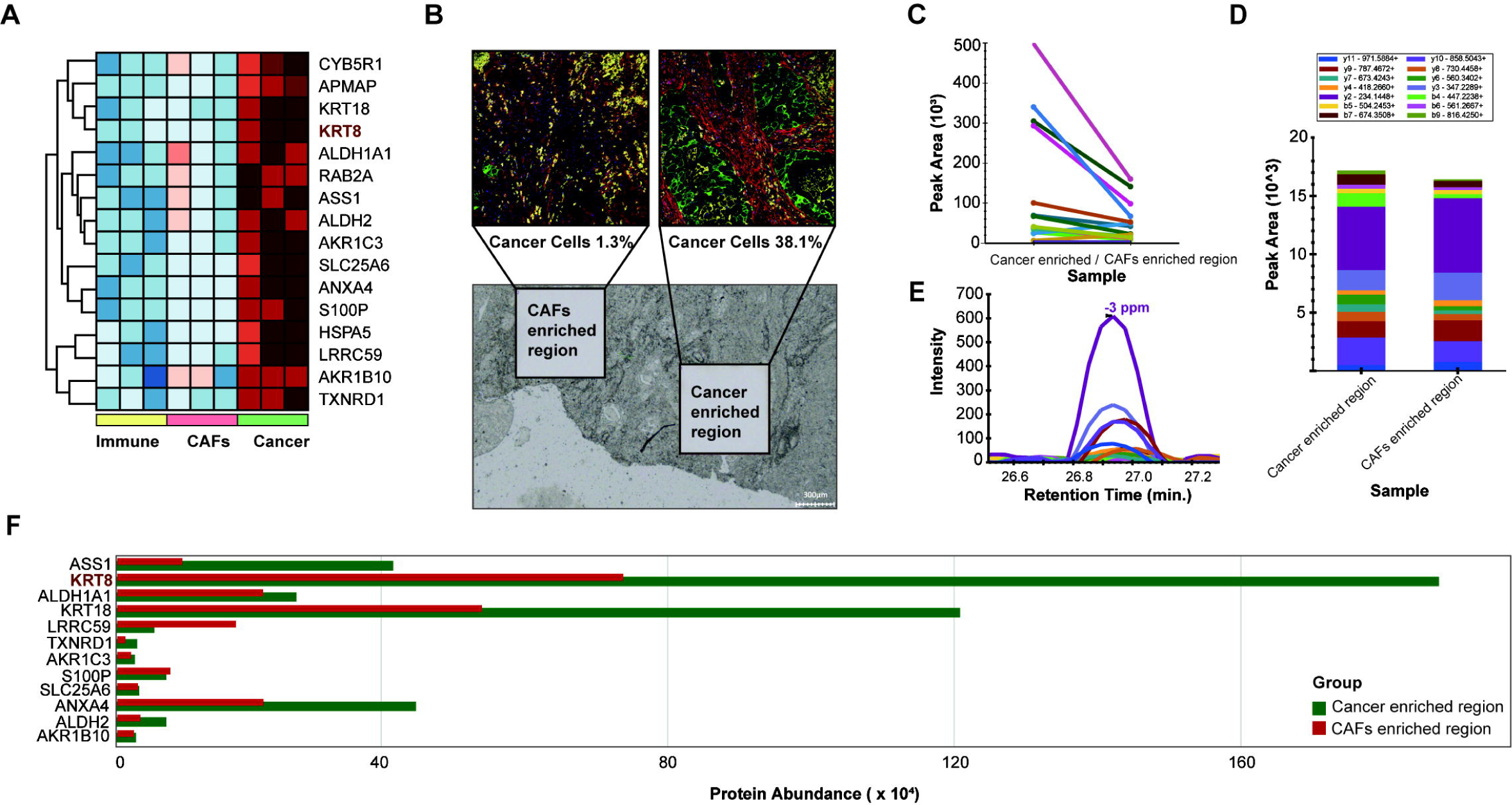
Targeted spatial proteomics. **(A)** Selection of cluster proteins with gastric cancer marker KRT8. **(B)** Selected ROIs for targeted spatial proteomics: CAFs-enriched region containing minimal cancer cells content of only 1.3% and Cancer-enriched region with cancer cells content of 38.1%. **(C)** Quantification of 14 peptides of KRT8. **(D)** Quantification of single peptide DFLAGGIAAAISK for protein SLC25A6 and its expression levels in Cancer enriched region and CAFs enriched region. **(E)** Chromatographic transition peak quality of the of the only peptide DFLAGGIAAAISK for protein SLC25A6. **(F)** Protein abundance of 12 proteins in Cancer enriched region and CAFs enriched region.

## Discussion

LC-MS/MS-based spatial proteomics, when combined with laser-capture microdissection, enables selective enrichment of specific tissue regions (29); the integration of mIHC imaging at cell-type resolution further allows precise isolation of marker-positive areas (6). Tools such as advanced MS acquisition modes and alignment software for LCM remain insufficiently evaluated. Here, we evaluated the Zeno TOF 7600 coupled with nanoLC as a practical platform. With standard K562 digest, Zeno SWATH identified over 7,000 protein groups from nanogram-scale samples. Notably, with sample inputs below 10 ng, nano-LC detected 4 to 5 times more precursors than the leading report (9). Moreover, ZT Scan mode improved reproducibility, with 45.7% of proteins showing a CV below 10%, thus mitigating variability in spatially restricted samples. Targeted analysis quantified 184 peptides per run, with sufficient data points per peak to support potential single-shot analysis of over 300 peptides. For spatial cell-type resolved proteomics in clinical FFPE tissues, we employed image-guided LCM with an optimized DIA workflow to compare cancer cells, CAFs, and immune cells. As a result, KRT8 was enriched in tumor cells, while stromal regions showed higher levels of extracellular matrix proteins. Collectively, these results demonstrated the robustness of the Zeno TOF 7600 platform for nanoLC-based proteomics, especially for spatial visual proteomics.

## Supplementary Data

Supplementary Figure S1-6

## Data Availability

The mass spectrometry proteomics data have been deposited to the ProteomeXchange Consortium via the PRIDE (30) partner repository with the dataset identifier PXD063580.

## Acknowledgments

We thank AB SCIEX for providing access to the Zeno TOF 7600 system with a Waters M-class HPLC, and for all the technical support provided during this study.

## Funding and additional information

China State Key Basic Research Program Grants (2024YFA1307200, 2021YFA1301601, 2020YFE0202200, 2022YFC3401104, 2021YFA1301602 and 2021YFA1302603), the National Natural Science Foundation of China (92253304, 22125403, 32201218, and 22104047), the Shenzhen Innovation of Science and Technology Commission (JSGGZD20220822095200001, JCYJ20200109141212325, JCYJ20210324120210029 and JCYJ20200109140814408). Research Grants Council, Hong Kong SAR Government (Grant No. C5005-23W). China Postdoctoral Science Foundation (2021M701410), Shenzhen Medical Research Fund (2401008).

## Author contributions

R.T. and W.G conceived the original idea and led the conceptualization of the research project.

K.Z. performed the data collection, conducted formal analysis, and prepared the original draft of the manuscript. B.Z. was responsible for sample processing and instrument maintenance. J.W. conducted comprehensive image analysis and figure preparation. A.H. contributed to data visualization. X.W. carried out the investigation and experimental procedures. H.W. and Q.L provided and characterized clinical samples. H.L and R.X supervised the project, reviewed the methodology, and provided critical guidance throughout the study.

## Declaration of Generative AI and AI-Assisted Technologies in the Writing Process

In the preparation of this manuscript, ChatGPT 4o was utilized to assist with language refinement, drafting, or editing suggestions. The authors take full responsibility for the content of the manuscript and have thoroughly reviewed and edited all outputs generated by the AI tool to ensure their accuracy, alignment with academic standards, and contextual appropriateness.

## Highlights

- Evaluation of the ZenoTOF 7600 with nano-LC ensuring sensitivity and reproducibility.
- Image-based segmentation and navigation for mIHC FFPE tissue LCM.
- Cell-type-specific spatial proteomic profiling with improved efficiency.
- Targeted proteomics for spatially resolved protein analysis within the same workflow.

