## Supplemental Figures for "Evaluation of the nano Liquid Chromatography coupled with Zeno TOF 7600 Mass Spectrometer for Cell Type-Resolved Spatial Proteomics"

**Highlights**

- Evaluation of the ZenoTOF 7600 with nano-LC ensuring sensitivity and reproducibility.
- AI-based segmentation and navigation for mIHC FFPE tissue analysis.
- Cell-type-specific spatial proteomic profiling with shortened gradients to improve efficiency.
- Targeted quantification integrated for spatially resolved protein analysis within the same workflow.

**Supplemental Figures**

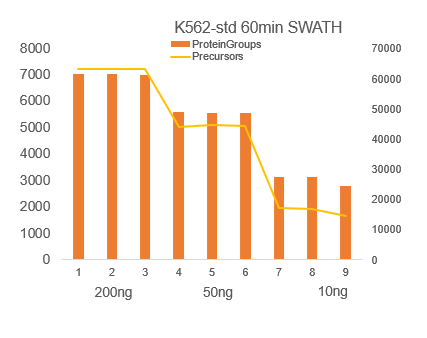

Figure S1A Identification number of zenoSWATH 20-200ng K562 without summarizing

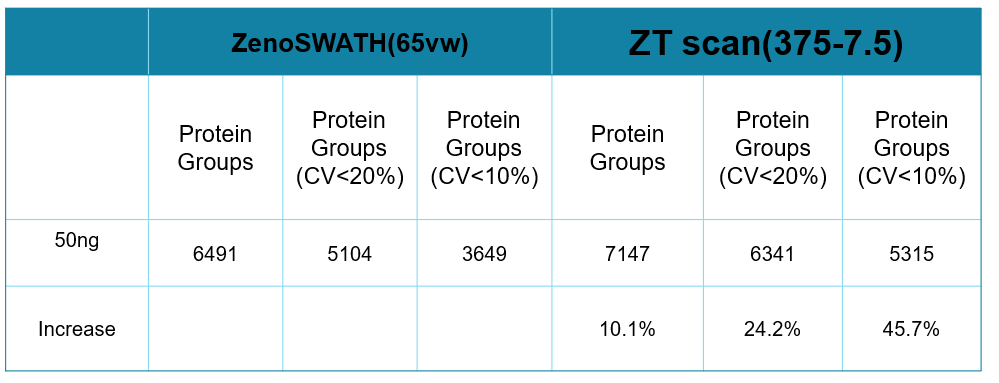

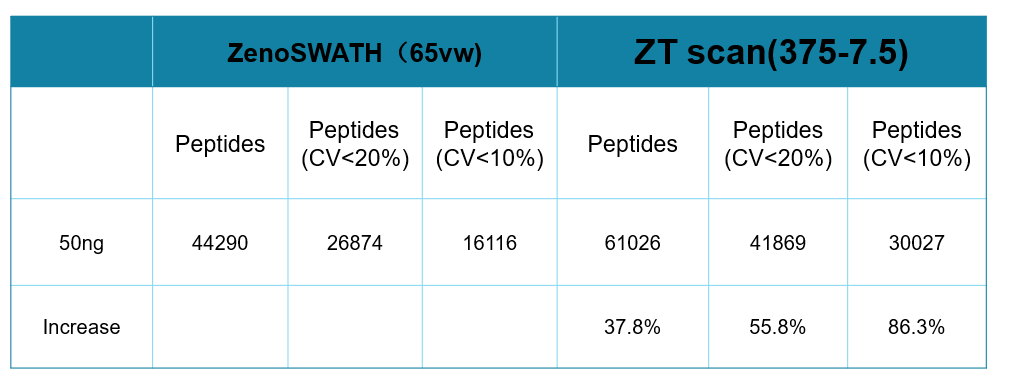

Figure S1B zenoSWATH and ZTscan comparison by 50ng K562

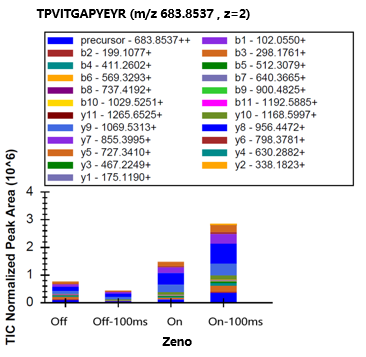

Figure S2A zeno 10ms and 100ms accumulation method on and off

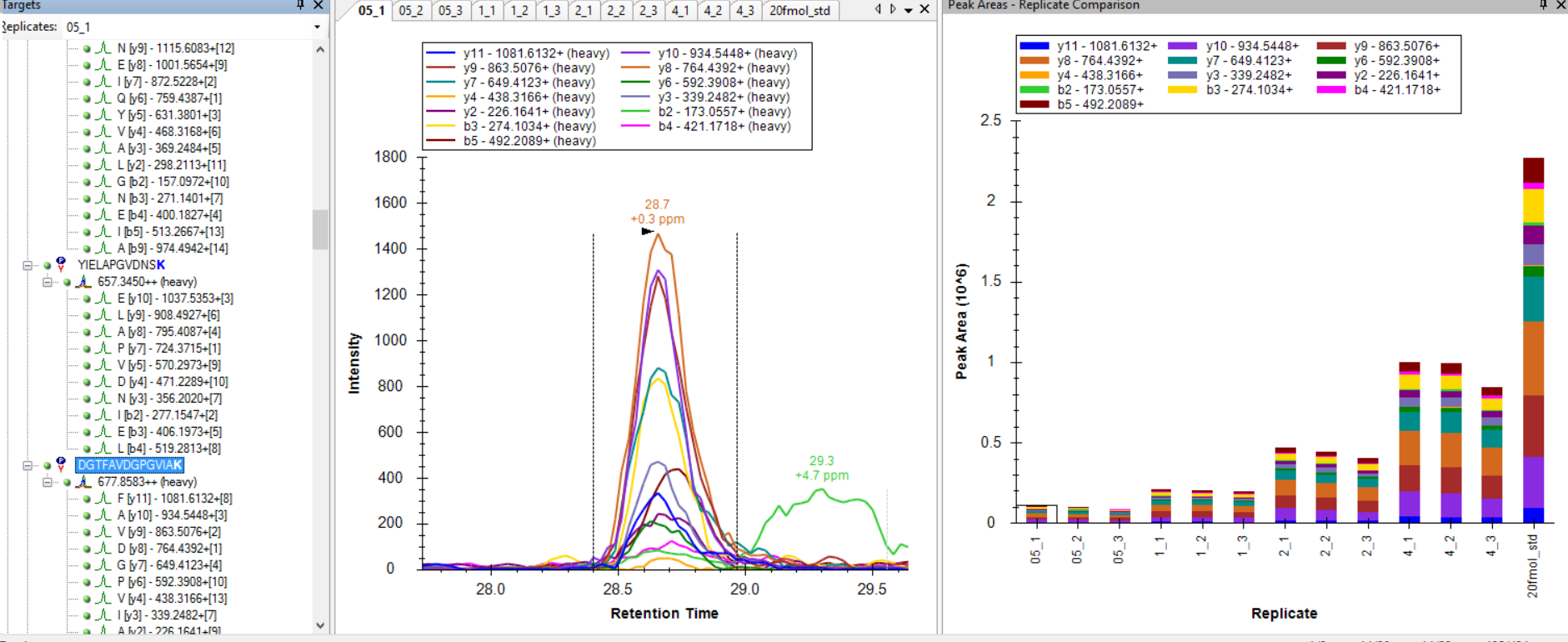

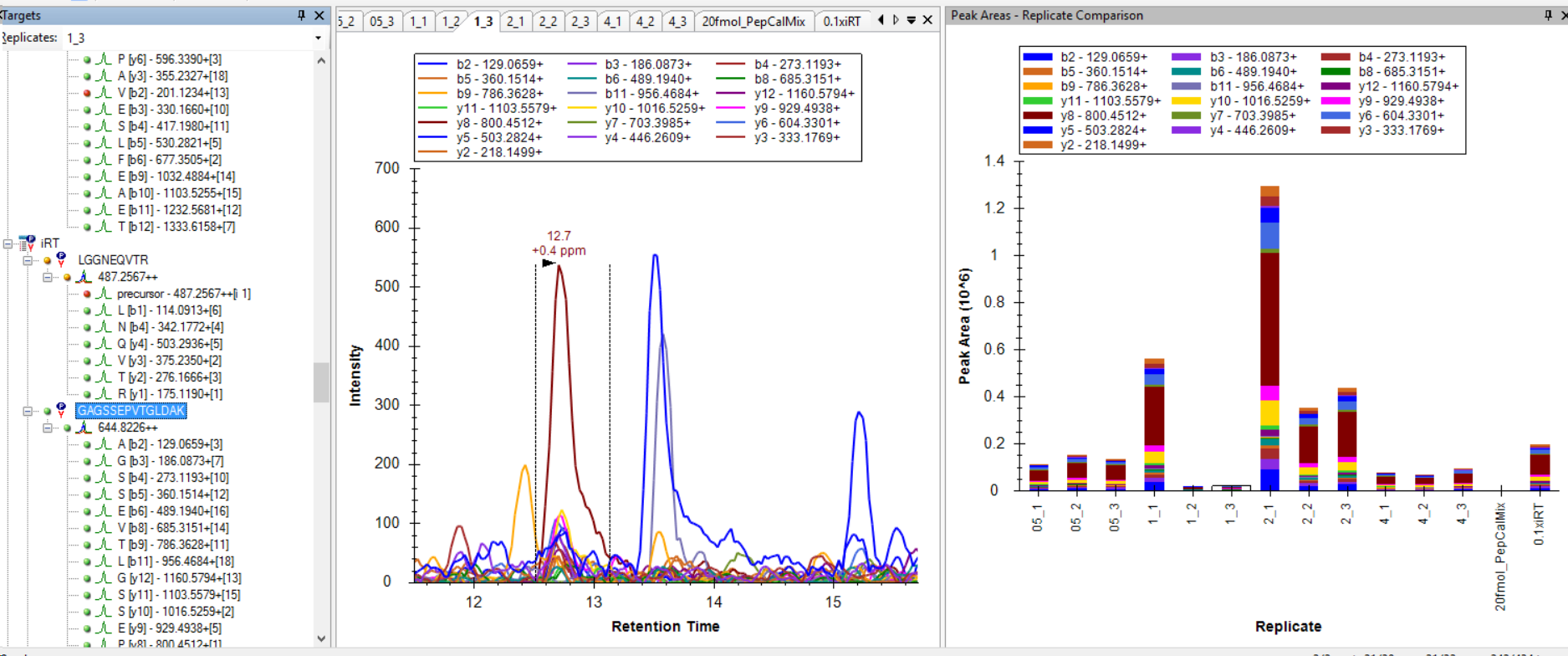

Figure S3 peptide DGTFAVDGPGVIAK peak quality and intensity and peptide GAGSSEPVTGLDAK peak quality and intensity

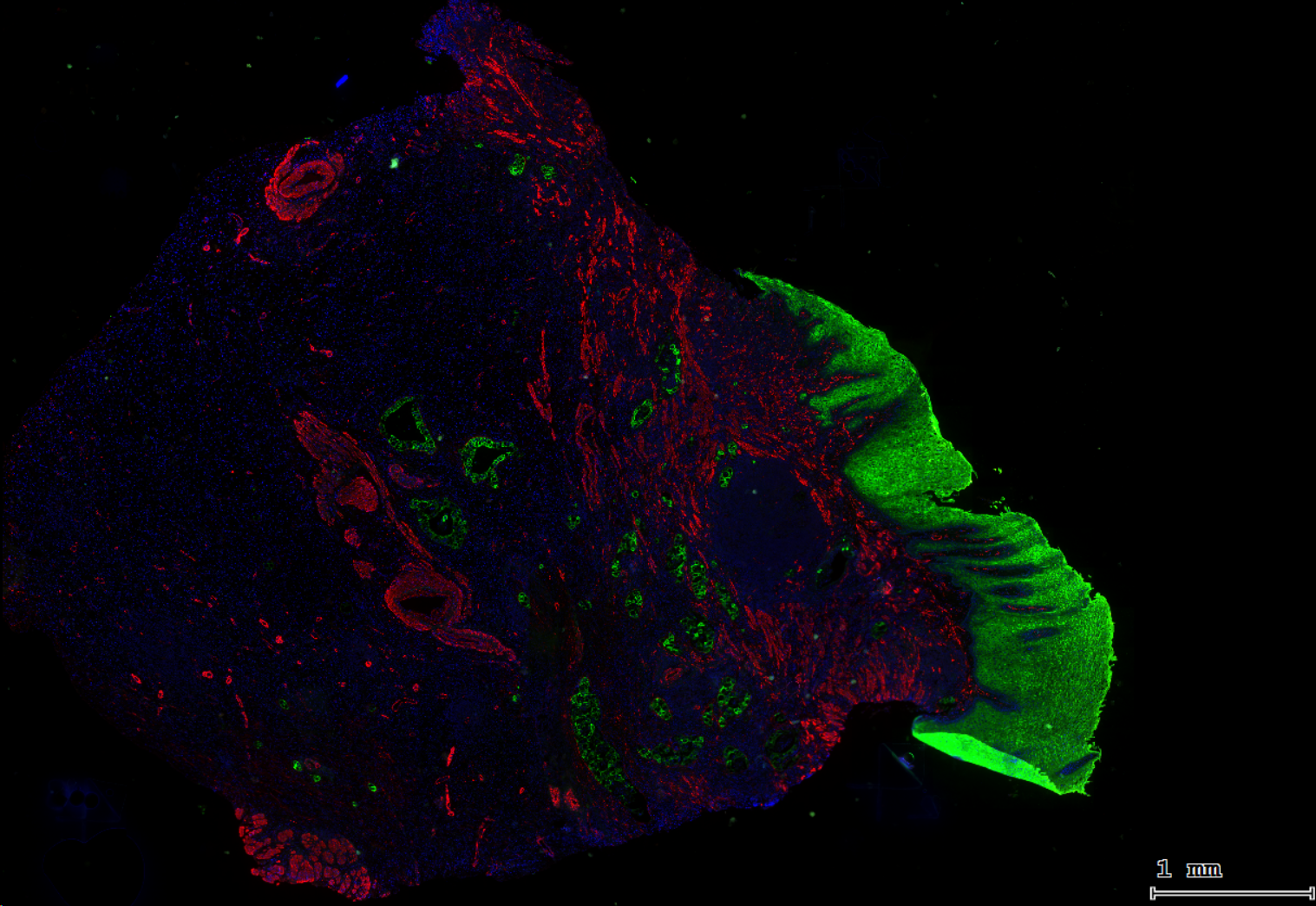

Figure S4 mIHC staning with ZT18-2 sample

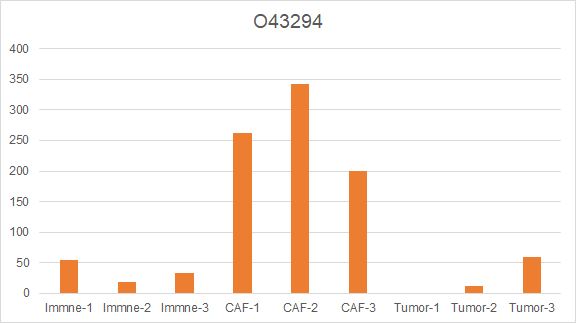

Figure S5 PG.Quantity of TGFB1 original value of spectronuant result

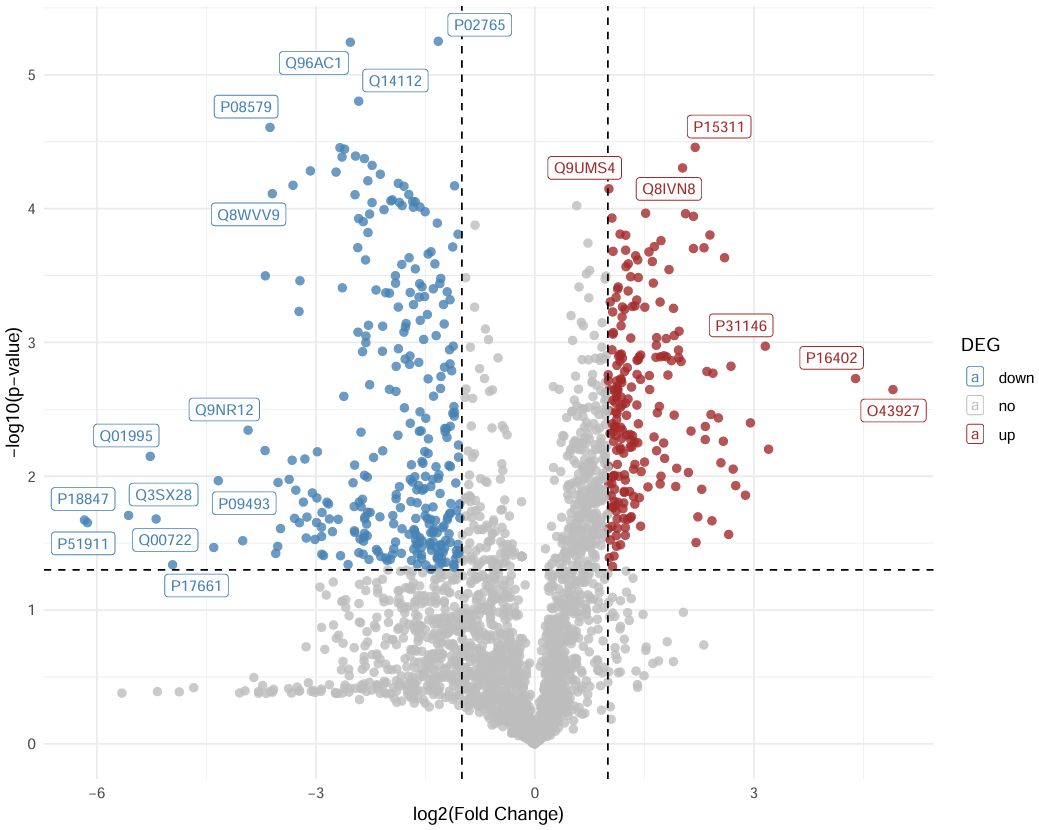

Figure S6 Volcano plot of CAF vs Immune groups
